# Low dose AKT inhibitor miransertib cures PI3K-related vascular malformations in preclinical models of human disease

**DOI:** 10.1101/2021.07.16.452617

**Authors:** Piotr Kobialka, Helena Sabata, Odena Vilalta, Ana Angulo-Urarte, Laia Muixí, Jasmina Zanoncello, Oscar Muñoz-Aznar, Nagore G. Olaciregui, Cinzia Lavarino, Veronica Celis, Carlota Rovira, Susana López, Eulàlia Baselga, Jaume Mora, Sandra D. Castillo, Mariona Graupera

## Abstract

Low-flow vascular malformations are congenital overgrowths composed by abnormal blood vessels potentially causing pain, bleeding, and obstruction of different organs. These diseases are caused by oncogenic mutations in the endothelium which result in overactivation of the PI3K/AKT pathway. Lack of robust *in vivo* preclinical data has prevented the development and translation into clinical trials of specific molecular therapies for these diseases. Here, we describe a new reproducible preclinical *in vivo* model of PI3K-driven vascular malformations using the postnatal mouse retina. This model reproduces human disease with *Pik3ca* activating mutations expressed in a mosaic pattern and vascular malformations formed in veins and capillaries. We show that active angiogenesis is required for the pathogenesis of vascular malformations caused by activating *Pik3ca* mutations. Using this model, we demonstrate that low doses of the AKT inhibitor miransertib both prevents and induces the regression of PI3K-driven vascular malformations. We confirmed miransertib efficacy in isolated human endothelial cells with genotypes spanning most of human low-flow vascular malformations.

**Graphical abstract:** 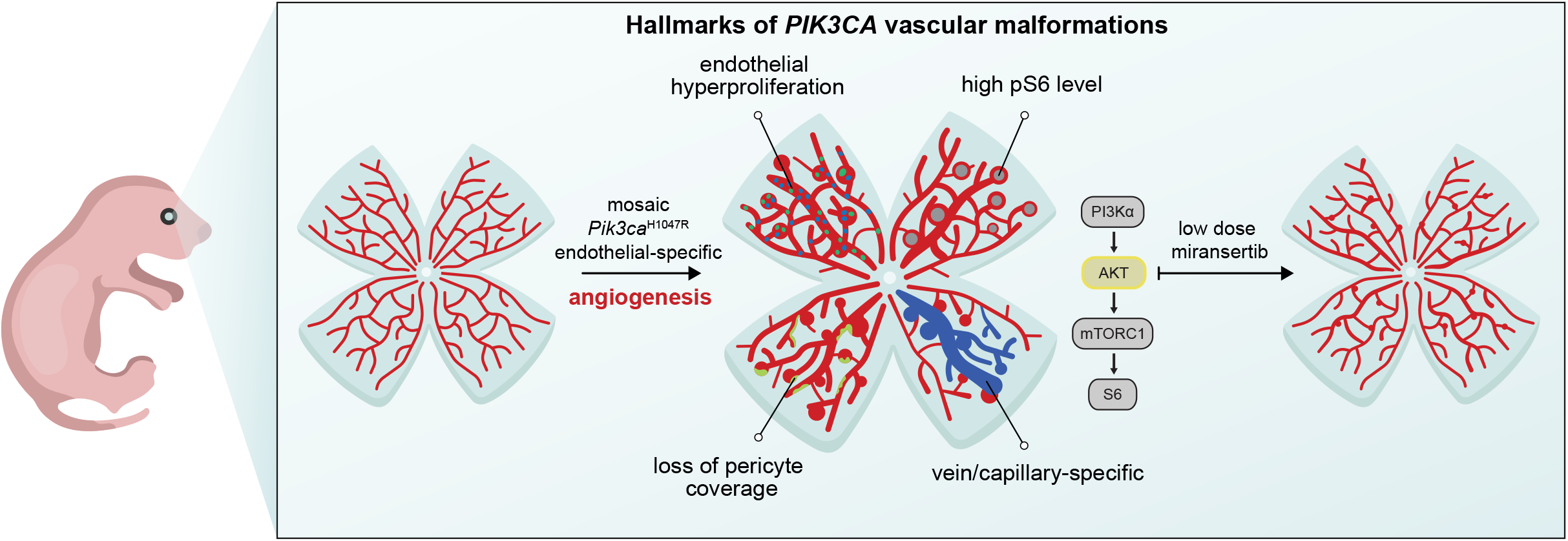

Low-flow vascular malformations are caused by PI3K signalling overactivation in endothelial cells. We have generated an optimised and robust preclinical system of PI3K-driven vascular malformations by inducing the mosaic expression of *Pik3ca*^*H1047R*^ in the retinal angiogenic endothelium. This preclinical model displays traits constituting the main hallmarks of the pathogenesis of low-flow blood vascular malformations: overactivation of PI3K signalling (high phospho-S6), vascular compartment specificity, loss of pericyte coverage, and endothelial cell hyperproliferation. Using this preclinical model we report that low dose AKT inhibitor miransertib prevents and regress PI3K-driven vascular malformations.

## Introduction

Vascular malformations are a congenital group of diseases composed of abnormal vascular channels that can occur anywhere in the body and potentially had a major impact on the quality of life of patients. They tend to be painful and disfiguring, and many leading to bleeding, recurrent infections, thrombosis, organ dysfunction, and even death (Van Damme *et al*, 2020). Vascular malformations can be classified as low-flow (venous, lymphatic and capillary) and fast-flow (arteriovenous) lesions, the former being the most frequent subtype. Vascular malformations appear during embryonic development, when vascular growth factors are produced at high levels, and expand proportionally with the physiological growth of the patient (Pang *et al*, 2020). Of note, vascular malformations may occur in isolation or as a part of a syndrome (Canaud *et al*, 2021). At present, there is no molecularly targeted therapy in the current management for these diseases. Instead, standard of care includes a broad spectra of mostly inefficient and invasive techniques including bandage compression, surgical excision, and sclerosing approaches (Castillo *et al*, 2019).

Overactivation of phosphoinositide 3-kinase (PI3K) signalling is a hallmark of most low-flow vascular malformations (Castillo *et al*, 2019; Mäkinen *et al*, 2021; Canaud *et al*, 2021). Sporadic venous malformations, the most common type of vascular malformations, are caused by gain-of-function mutations either in the endothelial tyrosine kinase receptor TEK/TIE2 or in the PI3K catalytic subunit alpha *PIK3CA* (Limaye *et al*, 2015; Castillo *et al*, 2016a; Castel *et al*, 2016); with *TEK* and *PIK3CA* mutations being mutually exclusive. Also, 80% of lymphatic malformations are caused by activating *PIK3CA* mutations (Boscolo *et al*, 2015; Luks *et al*, 2015; Mäkinen *et al*, 2021). In addition, venous, lymphatic and/or capillary malformations are frequently present in overgrowth syndromes caused by *PIK3CA* mutations, the so-called PROS (PIK3CA-related overgrowth spectrum) (Keppler-Noreuil *et al*, 2015). *PIK3CA* mutations in vascular malformations are similar to those found in epithelial cancer, being the missense mutations in the helical (*PIK3CA*^*E542K*^, *PIK3CA*^*E545K*^) and the kinase (*PIK3CA*^*H1047R*^) domains the most prevalent (Samuels *et al*, 2004).

*PIK3CA* encodes the p110α lipid kinase protein, which is a major signalling component downstream of growth factor receptor tyrosine kinases (RTKs) (Bilanges *et al*, 2019; Kobialka & Graupera, 2019). Specifically in endothelial cells, p110α is activated by the vascular endothelial growth factor receptors (VEGF-R) and TIE tyrosine kinase receptors (Graupera & Potente, 2013). Hence, it is not surprising that p110α is the sole class I PI3K isoform required for blood and lymphatic vascular development (Graupera *et al*, 2008; Stanczuk *et al*, 2015). p110α catalyses the phosphorylation of the lipid second messenger phosphatidylinositol-4,5-triphosphate (PIP_2_) to phosphatidylinositol-3,4,5-triphosphate (PIP_3_) at the cell membrane (Bilanges *et al*, 2019). PIP_3_, in turn, contributes to the recruitment and activation of a wide range of downstream targets, the serine-threonine protein kinase AKT (also known as protein kinase B, PKB) being critical in this cascade. The PI3K-AKT signalling pathway regulates many cellular processes that are key for endothelial cell biology, including cell proliferation, survival, and motility (Bilanges *et al*, 2019; Manning & Toker, 2017). There are three isoforms of AKT (AKT1, 2 and 3), showing high homology but being not redundant. AKT1 and AKT2 are broadly expressed, with AKT1 being the predominant isoform in endothelial cells (Chen *et al*, 2005; Ackah *et al*, 2005). *PIK3CA* and *TEK* gain-of-function mutations in endothelial cells lead to AKT hyper-phosphorylation which result in enhanced endothelial cell proliferation and resistance to cell death induced by growth factor withdrawal (Cai *et al*, 2019; Le Cras *et al*, 2020). Of note, pathological proliferation burst of cultured *PIK3CA* and *TEK* mutant cells is elicited by supplementation with external growth factor signals.

The discovery that most low-flow lesions are caused by overactivation of PI3K signalling has catalysed the repurpose of PI3K pathway inhibitors for these diseases. Given that pathological endothelial mutant cells primarily depend on AKT signalling, inhibition of AKT is a promising strategy for low-flow vascular malformations. Amongst AKT inhibitors, miransertib (ARQ 092, MK-7075) is a potent and selective allosteric AKT inhibitor showing higher specificity for the AKT1 isoform. Miransertib suppresses AKT activity by inhibiting membrane-bound active form of AKT and preventing activation of the inactive form of AKT (Yu *et al*, 2015). This inhibitor has shown efficacy in preclinical studies for PI3K-driven tumours (Yu *et al*, 2015, 2017) and Proteus syndrome (Lindhurst *et al*, 2015), which is caused by a somatic AKT gain-of-function mutation. Also, compassionate use of this inhibitor has shown therapeutic efficacy in patients with Proteus and PROS (Leoni *et al*, 2019; Biesecker *et al*, 2020; Forde *et al*, 2021).

Here, we show that active angiogenesis is required for the formation of *Pik3ca*-driven vascular malformations and report a unique *in vivo* model of PI3K-driven vascular malformations that allows a more accurate understanding of the dynamic pathogenesis of these diseases and thus a more efficient assessment of therapeutic strategies. In addition, we show how analysing patient-derived endothelial cells from vascular malformations allows for personalised medicine testing in these diseases. Using a spectra of preclinical models we demonstrate the efficacy of the AKT inhibitor miransertib at low dose for PI3K-driven vascular malformations both for prevention and treatment strategies. Furthermore, we show that endothelial cells from *TEK* and *PIK3CA*-mutant vascular malformations respond similarly to miransertib.

## Results

### Active angiogenesis is required for the formation of PI3K-driven vascular malformations

Because vascular malformations appear during embryonic development when endothelial mitogenic signals are produced at high concentration, we postulated that the genesis of these lesions depends on the stage of vascular development. To test this idea, we took advantge of the mouse retinal vasculature which allows the study of the different phases of angiogenesis. We chose three developmental stages (early, intermediate and late) in which endothelial cells (ECs) exhibit different dependency to mitogenic signals (Ehling *et al*, 2013). We induced the endogenous expression of the *Pik3ca*^H1047R^ mutation (Kinross *et al*, 2012) in heterozygosis by using the Pdgfb-iCreER mice, which expresses a tamoxifen-inducible Cre recombinase specifically in ECs (Claxton *et al*, 2008). 4-Hydroxytamoxifen (4-OHT) was administered at postnatal day (P)1, P7 or P15 and retinas were isolated one week later (Fig 1A). By analysing the extent of vascular overgrowth, we found that both early and intermediate stages showed full penetrance, with all retinas analysed developing vascular malformations (Fig 1B,C). However, the degree of enhanced vascularity was more prominent and generalized in the early develomnetal stage than in the intermediate period (Fig 1B-D). In contrast, only one third of the retinas showed malformed vascular areas when *Pik3ca*^H1047R^ mutation was expressed at P15 (Fig 1B-D). By taking advantage of ROSA^mTmG^;Pdgfb-iCreER (later referred to as EC-mTmG) reporter mice (Muzumdar *et al*, 2007), which expresses cell membrane-localized EGFP following Cre recombination, we demonstrated that the Pdgfb-iCreER line is similarly active at all time points tested (Fig 1E, F). Thus, penetrance and severity of vascular malformations in our model are independent of the number of ECs recombined at the different stages. These results indicate that the expression of mutant *Pik3ca* in ECs is not sufficient for the acquisition of a malformed vascular phenotype and that active angiogenesis is required for PI3K-driven vascular malformations to occur.

**Figure 1.**
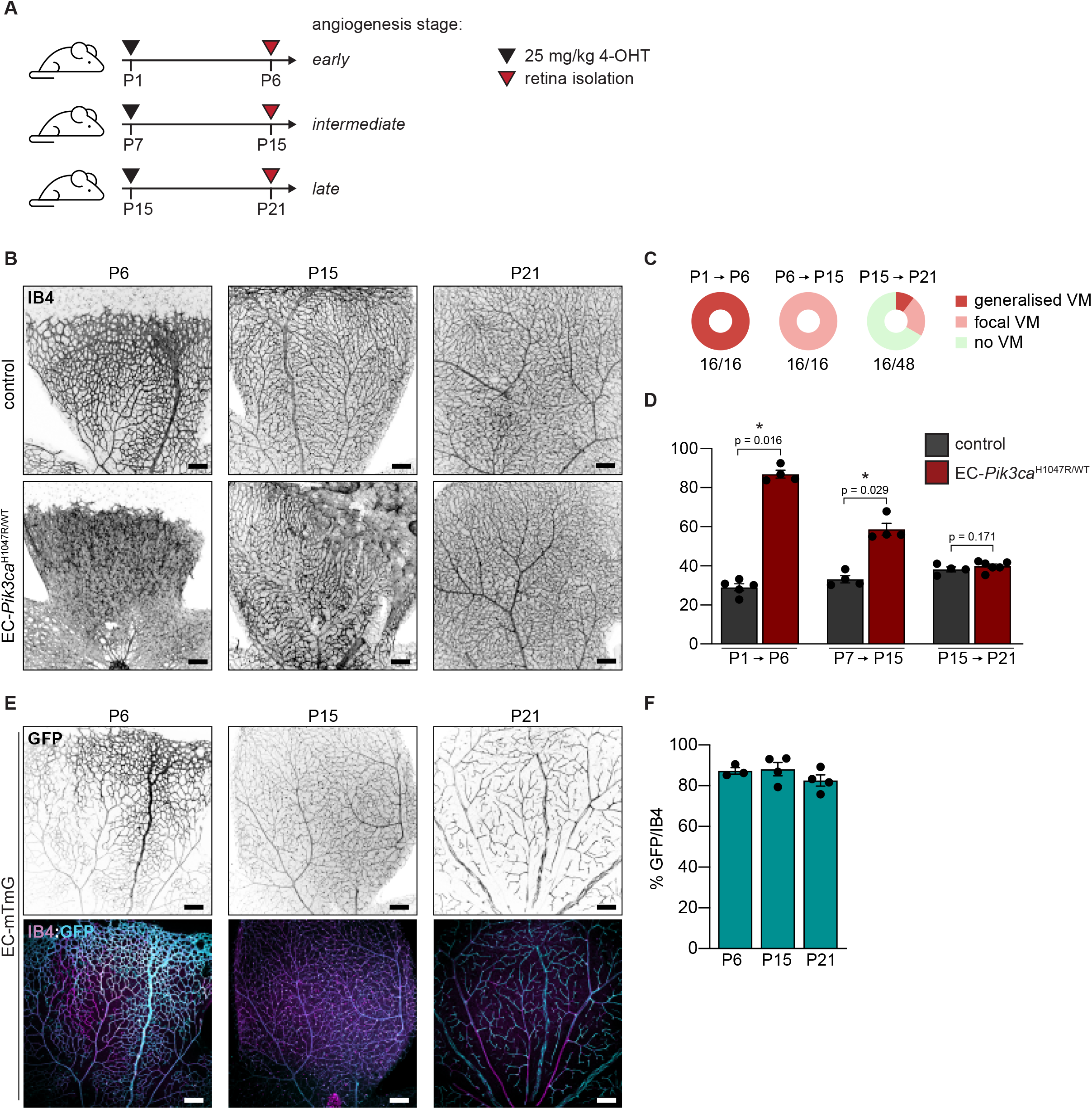
Pathogenesis of *Pik3ca*-driven vascular malformations depends on active angiogenesis. (**A**) Experimental setup scheme showing analysed angiogenic stages in the retina models. (**B**) Representative images of control and EC-*Pik3ca*^H1047R/WT^ mouse retinas isolated at indicated time points stained with IB4 for blood vessels. Scale bars: 150 μm. (**C**) Pie charts showing the incidence of vascular lesions at different angiogenic stages. Lesions were categorised according to their expanse (generalised or focal). Quantification was performed per retina petal. Numbers below show the presence of any type of lesions per retina petal. (**D**) Quantification of IB4-positive areas per retina. Error bars are s.e.m. n ≥ 3 retinas per genotype. (**E**) Representative images of EC-mTmG mouse retinas isolated at indicated time points stained with IB4 and GFP. Scale bars: 150 μm. (**F**) Quantification of GFP/IB4 ratio per retina. Error bars are s.e.m. n ≥ 3 retinas per genotype. Statistical analysis was performed by nonparametric Mann– Whitney test. *p < 0.05 was considered statistically significant.

### A new preclinical model of PI3K-driven vascular malformations

Our data show that expression of *Pik3ca*^H1047R^ in ECs at an early stage of postnatal angiogenesis leads to generalised vascular malformations. However, sporadic vascular malformations appear as a mosaic disease, where malformed vascular lesions tend to be isolated and focal. Hence, to better reproduce the etiology of human disease, we studied the impact of a decreasing range of 4-OHT doses during early developmental stage in EC-*Pik3ca*^WT/H1047R^ mouse retinas (Fig 2A). To validate mosaicism and identify targeted ECs in our dosing strategy, we treated EC-mTmG P1 mice in parallel with the same doses (Fig 2B, C). This approach allowed us to identify the lowest 4-OHT dose (0.125 mg/kg) that led to distinguishable vascular malformations with a total vascular density significantly increased compared to wild type counterparts (Fig 2A, D). Also, we noticed that low dose of 4-OHT allowed the targeting of ECs of all vessel subtypes including arteries, veins, and capillaries (Fig 2B). Instead, expression of *Pik3ca*^H1047R^ upon low dose of 4-OHT resulted in the formation of vascular malformations only in veins and capillaries (Fig 3A). This is consistent with the observation that, within the blood vessel compartment, *PIK3CA* mutations are only present in human venous and capillary malformations (Keppler-Noreuil *et al*, 2015; Castillo *et al*, 2016a). *Pik3ca*^H1047R^-vascular malformations in postnatal retinas showed enriched phospho (p)-S6 (Ser235/236) levels, a read-out for PI3K/AKT/mTORC1 signalling, compared to the surrounding normal vasculature and wild type retinas (Fig 3B, D). This focal PI3K signalling activation resulted in EC hyperproliferation (Fig 3C, E) and accumulation of ECs, causing the overall enhanced vascularity (Fig 3C, F, G). These lesions also exhibited loss of pericyte coverage, assessed by immunostaining for pericyte-specific marker NG2, in contrast to non-malformed vasculature in the same retina and the control (Fig EV3). Collectively, these results show that EC-specific mosaic induction of endogenous *Pik3ca*^*H1047R*^ expression during active vascular growth faithfully models human low-flow vascular malformations.

**Figure 2.**
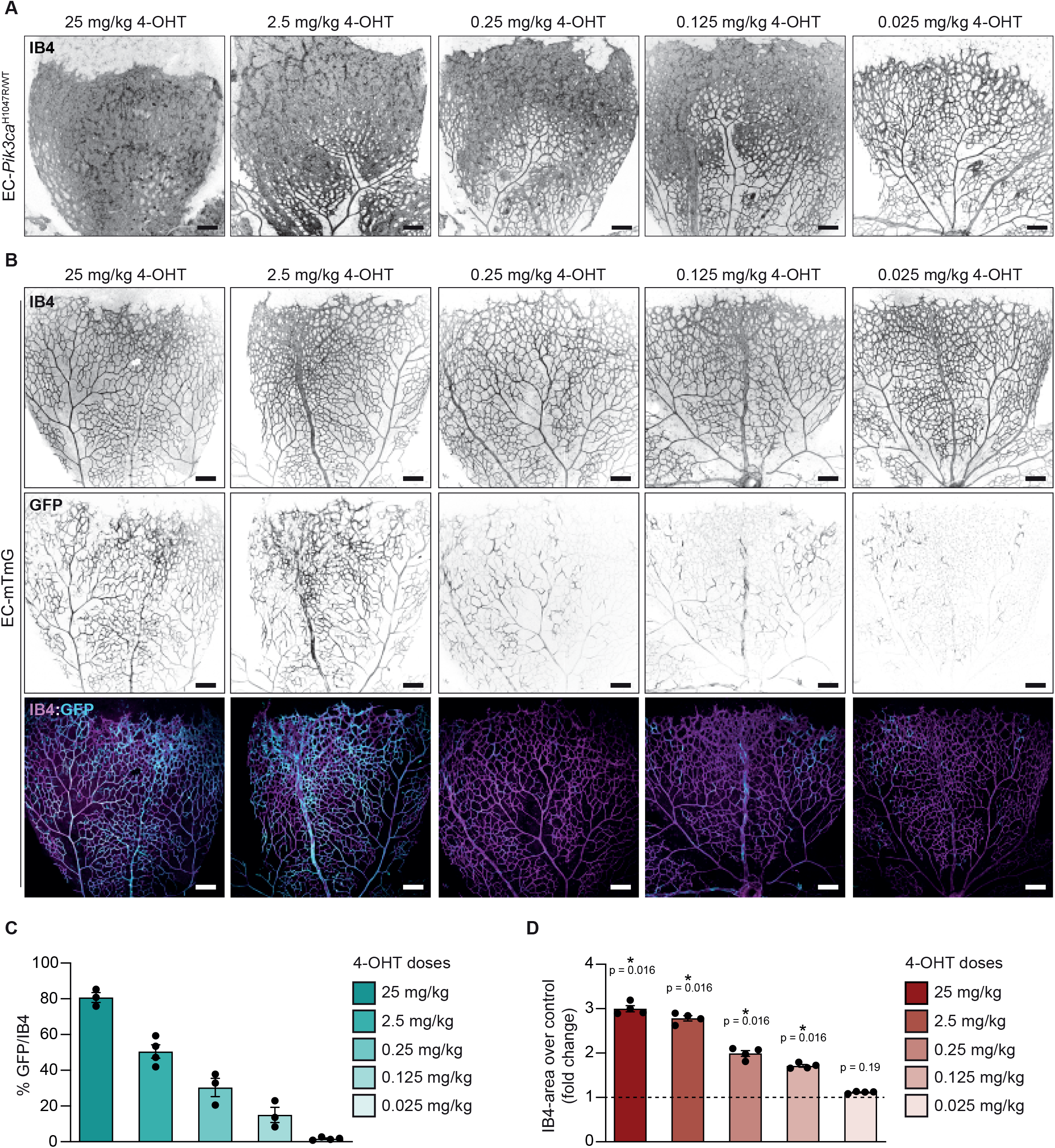
Modelling PI3K-vascular malformations in murine retinas by mosaic expression of Pik3ca^H1047R^ in ECs. (**A, B**) Representative images of EC-*Pik3ca*^H1047R/WT^ (**A**) and EC-mTmG (**B**) P6 retinas from mice treated with decreasing doses of 4-OHT on P1. Retinas were stained for blood vessels (IB4) and GFP as indicated. Scale bars: 150 μm. (**C**) Quantification of GFP/IB4 ratio of EC-mTmG retinas. Error bars are s.e.m. n ≥ 3 retinas per genotype. (**D**) Quantification of IB4-positive area per retina in *Pik3ca*^H1047R/WT^ retinas. Data presented as a percentage of the control for each 4-OHT dose. Error bars are s.e.m. n = 4 retinas per genotype. Statistical analysis was performed by nonparametric Mann–Whitney test. *p < 0.05 was considered statistically significant.

**Figure 3.**
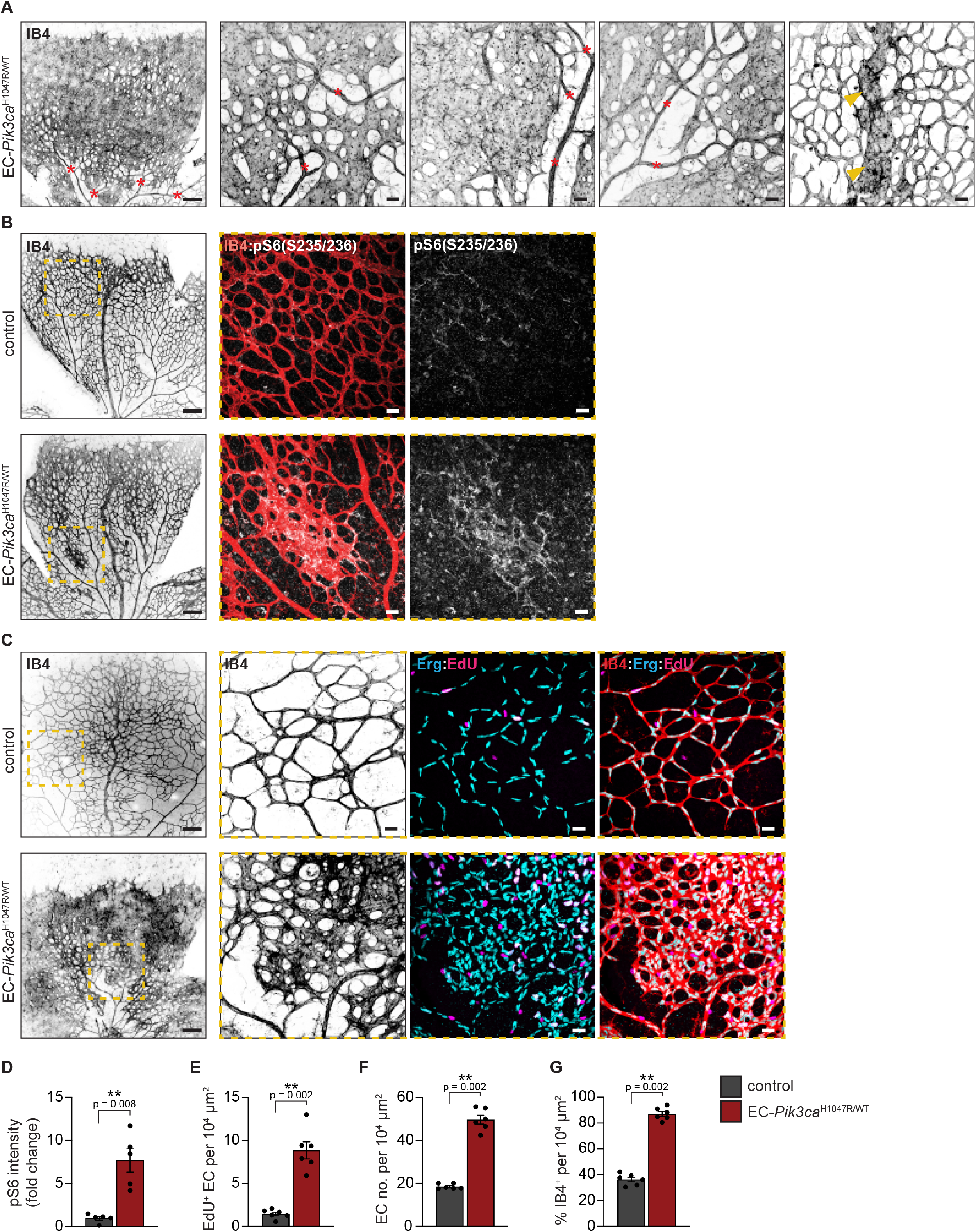
*Pik3ca*-vascular malformations in murine retinas reproduce hallmarks of human disease. (**A**) Representative images of EC-*Pik3ca*^H1047R/WT^ P6 retinas isolated from mice treated with 0.125 mg/kg of 4-OHT on P1. Retinas were immunostained for blood vessels (IB4). Red asterisks show arteries and arterioles and yellow arrowheads veins. Scale bars: 150 μm (left panel) and 30 μm (right, high magnification panels). (**B**) Representative images of P6 retinas from control and EC-*Pik3ca*^H1047R/WT^ mice treated with 0.125 mg/kg 4-OHT on P1, following immunostaining for p-S6 (S235/236) and blood vessels (IB4). Scale bars: 150 μm (left panels) and 30 μm (right panels, high magnification). (**C**) Representative control and EC-*Pik3ca*^H1047R/WT^ P6 retinas immunostained for blood vessels (IB4), EC nuclei (Erg) and EdU. Scale bars: 150 μm (left panels) and 30 μm (right panels, high magnification). Quantification of (**D**) p-S6 (S235/236) intensity (presented as a fold change of vehicle-treated control), (**E**) EC proliferation by EdU staining, (**F**) EC number by Erg positive cells and (**G**) retinal vascularity by IB4-positive area in control and EC-*Pik3ca*^H1047R/WT^ retinas. Error bars are s.e.m. n > 5 retinas per genotype. Statistical analysis was performed by nonparametric Mann–Whitney test. **p < 0.01 was considered statistically significant.

### Low dose AKT inhibitor miransertib prevents the formation of PI3K-driven vascular malformations

Targeting AKT, the main player of PI3K-driven signalling in ECs, has not yet been assessed *in vivo* in vascular malformations. Thus, we took advantage of our unique *in vivo* model to examine the impact of miransertib in the dynamic pathophysiology of PI3K-driven vascular malformations. Previous preclinical studies of miransertib using tumor xenografts showed that the minimum dose that has an impact on tumour volume is 75 mg/kg (Yu *et al*, 2017); thus, we first evaluated this dose to assess miransertib for preventing the formation of *Pik3ca*-driven vascular malformations. For this, we treated P1 EC-Pik3ca^WT/H1047R^ mice with 4-OHT and we dosed these mice with either 75 mg/kg of miransertib or vehicle at P1 and P2, followed by the analysis of P6 retinas (Fig 4A). Miransertib prevented the formation of vascular malformations, assessed by the vascular area and total EC number (Fig 4B, D, E) by inhibiting *Pik3ca*^H1047R^-driven EC hyperproliferation assessed by EdU incorporation (Fig 4B, F). Also, treatment prevented the loss of NG2-positive mural cell coverage (Fig EV4). By using pS6 immunostaining, we confirmed that miransertib treatment inhibited PI3K signalling in the vasculature (Fig 4C, G). Of note, wild type control retinas treated with miransertib showed a slight impact on vasculature density and EC proliferation (Fig 4B, D, F) further supporting a key role of AKT in angiogenesis and vascular homeostasis (Kerr *et al*, 2016; Ackah *et al*, 2005; Chen *et al*, 2005).

**Figure 4.**
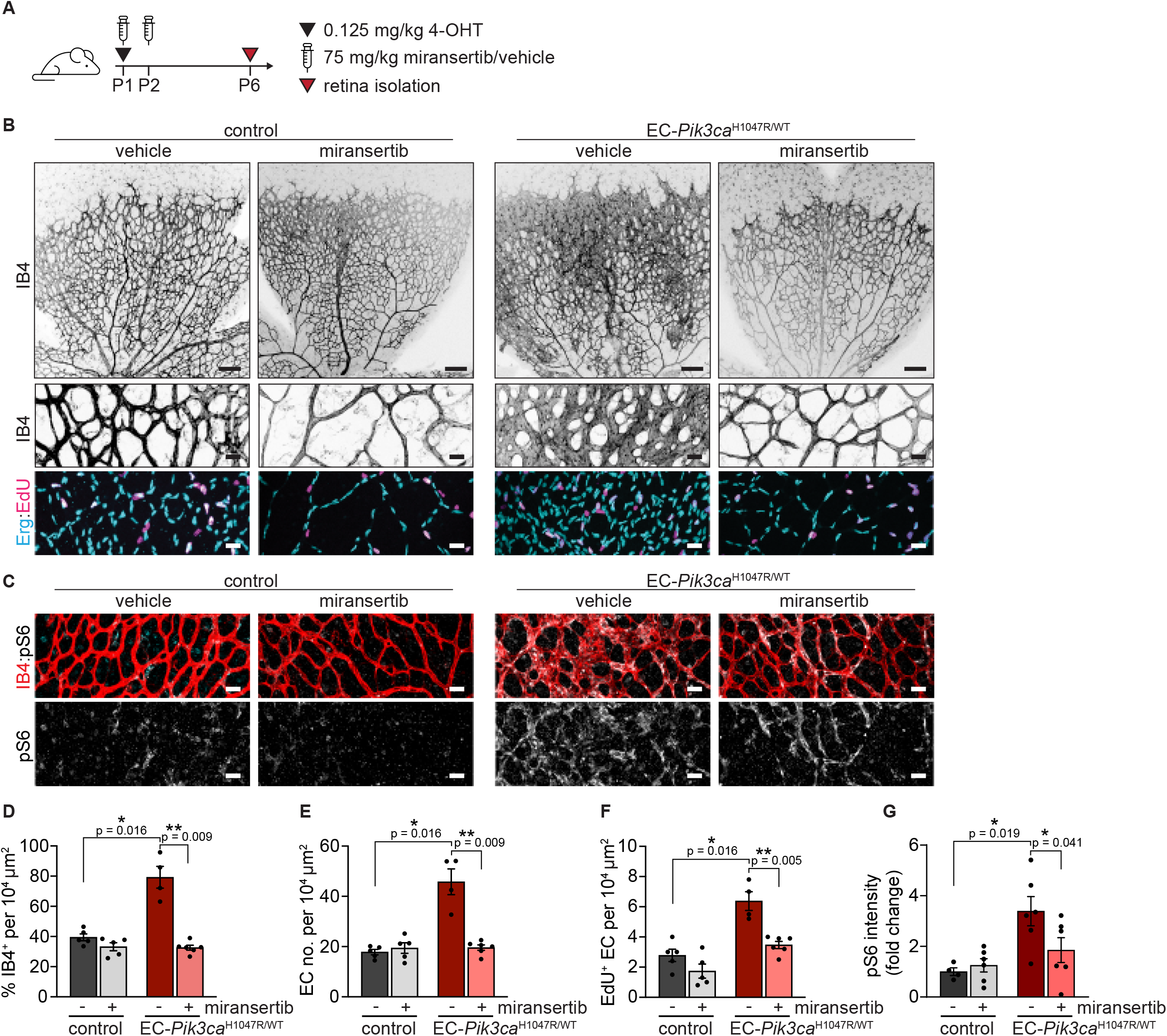
Miransertib prevents the formation of *Pik3ca*-vascular malformations in mice. (**A**) 4-OHT and miransertib dosing scheme used for a prevention therapeutic experimental setup. (**B, C**) Representative images of P6 retinas isolated from control and EC-*Pik3ca*^H1047R/WT^ mouse littermates. Blood vessels were stained with IB4. Lower panels showing high magnification images of the representative areas showing (**B**) blood vessels (IB4), EC nuclei (Erg), EdU incorporation and (**C**) pS6 (S235/236). Quantification of (**D**) retinal vascularity by IB4 staining, (**E**) EC number by Erg immunostaining, (**F**) EC proliferation by EdU staining, and (**G**) pS6 (S235/236) intensity (presented as a fold change of vehicle-treated control). Scale bars: 150 μm (upper panel) and 30 μm (lower panels). n ≥ 4 retinas per genotype. Statistical analysis was performed by nonparametric Mann–Whitney test. *p < 0.05 and **p < 0.01 were considered statistically significant.

Molecularly targeted treatment of vascular malformations might require long-term, even chronic, therapeutic approaches in paediatric patients. This points to the importance of identifying the minimal effective dose of any candidate treatment. Thus, we next asked whether reducing the dose of miransertib would provide similar efficacy and reduce knock-on effects on normal vasculature in our preclinical model. To test this, we reduced miransertib dose by half (35 mg/kg) and evaluated its efficacy on preventing *Pik3ca*-driven vascular malformations (Fig 5A). We observed that the therapeutic efficacy of AKT inhibition by miransertib was maintained at the lower dose, with decreased EC proliferation leading to reduced number of total ECs and vascular density in EC-*Pik3ca*^H1047R^ miransertib-treated retinas compared to vehicle-treated counterparts (Fig 5B, D-F). Low dose of miransertib efficiently inactivated PI3K signalling as shown by reduced pS6 levels (Fig 5C, G). Importantly, this minimum effective dose had no impact on non-mutant vasculature (Fig 5B-G). Altogether, our data demonstrate that miransertib efficiently inhibits the formation of PI3K-driven vascular malformations and that lower doses than reported for oncological purposes of miransertib appear equally effective for preventive strategies.

**Figure 5.**
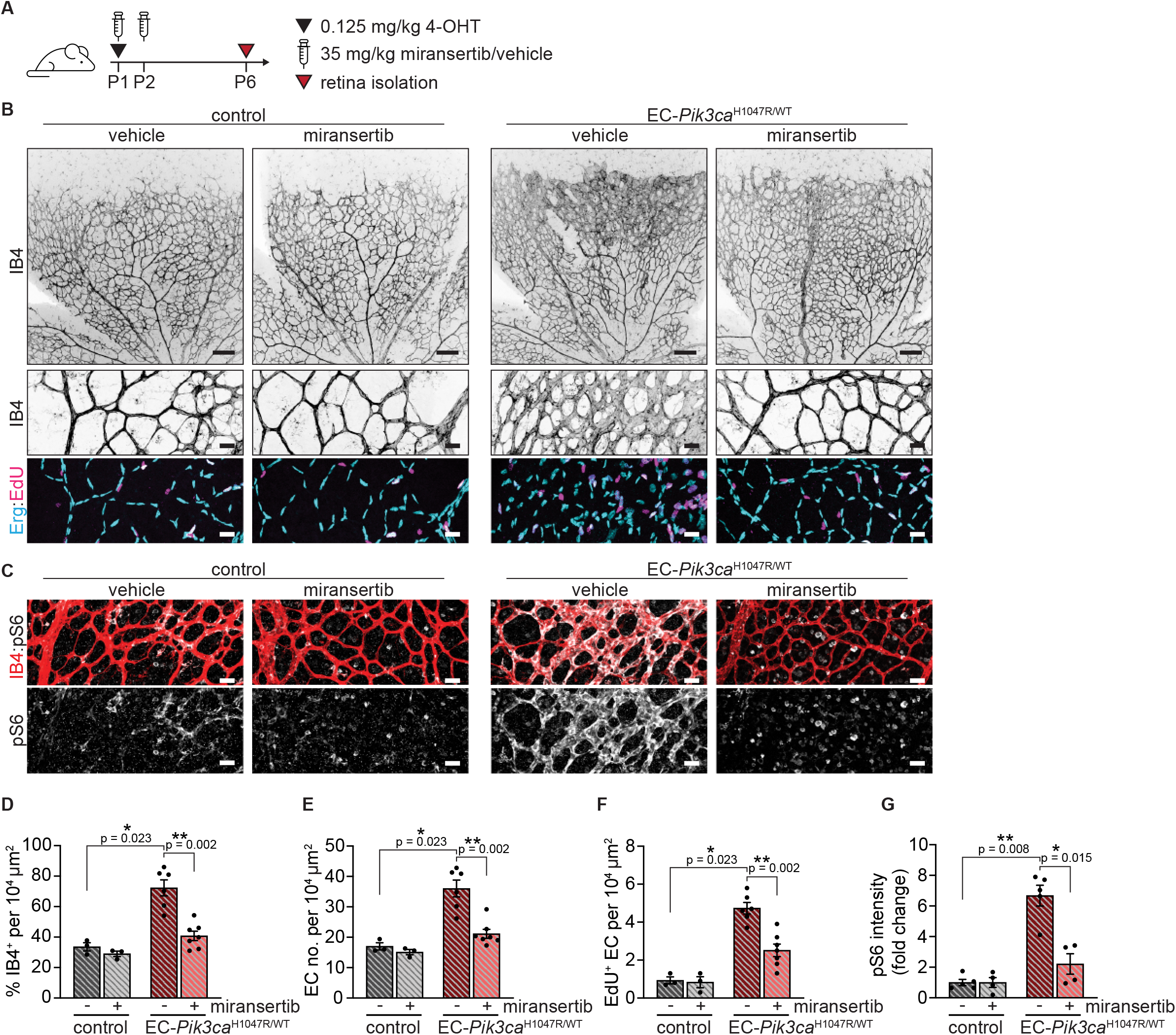
Low dose of miransertib prevents the growth of *Pik3ca*-vascular malformations. (**A**) 4-OHT and miransertib dosing scheme used for a prevention experimental setup. (**B, C**) Representative confocal images of P6 retinas isolated from control and EC-*Pik3ca*^H1047R/WT^ mouse littermates. Blood vessels were stained with IB4. Lower panels showing high magnification images of the representative areas showing (**B**) blood vessels (IB4), EC nuclei (Erg), EdU incorporation and (**C**) pS6 (S235/236). Quantification of (**D**) retinal vascularity by IB4 staining, (**E**) EC number by Erg immunostaining, (**F**) EC proliferation by EdU staining, and (**G**) pS6 (S235/236) intensity (presented as a fold change of vehicle-treated control). Scale bars: 150 μm (upper panels) and 30 μm (lower panels). n ≥ 3 retinas per genotype. Statistical analysis was performed by nonparametric Mann–Whitney test. *p < 0.05 and **p < 0.01 were considered statistically significant.

### Treatment with low dose of miransertib induces the regression of PI3K-driven vascular malformations

Since vascular malformations are congenital diseases, therapeutic interventions should aim for inducing the regression of mass lesions. In line with this, we assessed the therapeutic effect of low dose miransertib on PI3K-driven vascular malformations. For this, we first induced the formation of vascular malformations by treating EC-*Pik3ca*^WT/H1047R^ mice with 4-OHT at P1 and then treat these mice with 35 mg/kg (low dose) of miransertib at P4 and P5 (Fig 6A). Of note, in our *in vivo* preclinical model, vascular malformations are already present at P4 (Fig EV6), showing all hallmarks of PI3K-driven vascular malformations: increased vascularity (Fig EV6B, F), number of ECs (Fig EV6C, G), EC hyperproliferation (Fig EV6C, H), enhanced pS6 levels (Fig EV6D, I) and impaired pericyte coverage (Fig EV6E, J). In contrast to vehicle-treated, miransertib treatment of EC-*Pik3ca*^H1047R^ retinas effectively regressed vascular malformations, assessed by the normalisation of the vascular density, EC numbers, and proliferation rate (Fig 6B, D-F). Importantly, elevated PI3K signalling induced by *Pik3ca*^H1047R^ was eficiently blunted by miransertib (Fig 6C, G). These data demonstrate that low dose miransertib is effective in inducing regression of PI3K-driven vascular malformations; hence opening a very promising clinical strategy for these diseases.

**Figure 6.**
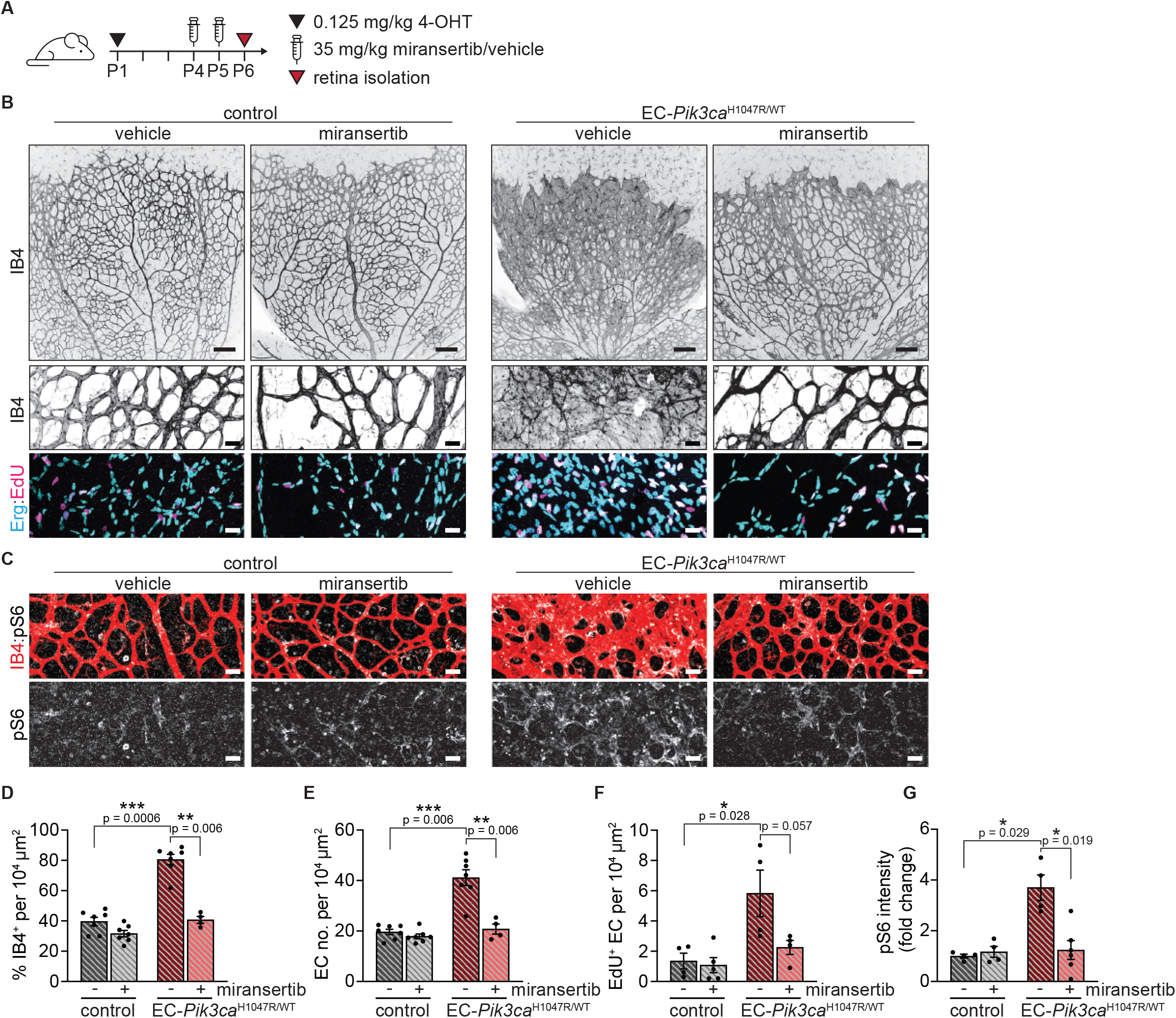
Low dose miransertib induces regression of *Pik3ca*-vascular malformations *in vivo*. (**A**) 4-OHT and miransertib dosing scheme used for a curative experimental setup. (**B, C**) Representative confocal images of P6 retinas isolated from control and EC-*Pik3ca*^H1047R/WT^ mouse littermates. Blood vessels were stained with IB4. Lower panels showing high magnification images of the representative areas showing (**B**) blood vessels (IB4), EC nuclei (Erg), EdU incorporation and (**C**) pS6 (S235/236). Quantification of (**D**) retinal vascularity by IB4 staining, (**E**) EC number by Erg immunostaining, (**F**) EC proliferation by EdU staining, and (**G**) pS6 (S235/236) intensity (presented as a fold change of vehicle-treated control). Scale bars: 150 μm (upper panels) and 30 μm (lower panels). n ≥ 4 retinas per genotype. Statistical analysis was performed by nonparametric Mann–Whitney test. *p < 0.05 and **p < 0.01 were considered statistically significant.

### Miransertib inhibits PI3K/AKT signalling and reduces cell viability in patient-derived PIK3CA- and TEK-mutant endothelial cells

Next, we assessed the therapeutic potential of miransertib in a preclinical human setting. For this, we isolated and cultured ECs derived from human vascular malformations carrying mutations in *PIK3CA* and *TEK/TIE2* (Appendix Table S1), spanning the genetic causes of more than 80% of low-flow lesions in patients (Castillo *et al*, 2016a; Castel *et al*, 2016; Limaye *et al*, 2015; Limaye *et al*, 2009). To isolate and culture these cells, fresh surgical resections of low-flow vascular malformations were subjected to tissue digestion and EC positive selection (Fig 7A). Sanger sequencing and droplet digital PCR revealed that causing mutations were present in the EC culture, with variant allelic frequencies (VAFs) of around 50% in these cell cultures (Fig 7B; Appendix Fig S1A,B). Characterization of these cells validated their specific EC properties – cobblestone morphology and expression of the EC-specific markers VE-cadherin and ERG (Fig 7C; Appendix Fig S1C). Next, we analysed the impact of these mutations in PI3K signalling by assessing AKT and S6 phosphorylation levels. Indeed, *PIK3CA*- and *TIE2*-mutant ECs exhibited constitutive activation of the pathway compared with wild type HUVECs (Fig 7D). To evaluate the therapeutic efficacy of miransertib in patient-derived *PIK3CA* and *TEK* mutant ECs, we first assessed its impact on PI3K/AKT signalling. At very low doses, miransertib strongly inhibited AKT signalling assessed by AKT and S6 phosphorylation levels that were reduced in a dose-dependent manner. This effect was similar in both *PIK3CA* and *TEK* mutant ECs (Fig 7E, F). We then studied the functional impact of miransertib-mediated AKT signalling inhibition by determining the dose-response effect on cell viability in ECs with either *PIK3CA*- or *TEK*-mutant genotype. Miransertib robustly decreased EC viability in both genotypes showing low IC_50_ with overlaping (not significantly different) confident intervals (Fig 7G). These data show that miransertib impacts the viability of patient-derived ECs at low concentrations and that it might constitute a promising therapeutic strategy for both *PIK3CA* and *TEK*-mutant vascular malformations.

**Figure 7.**
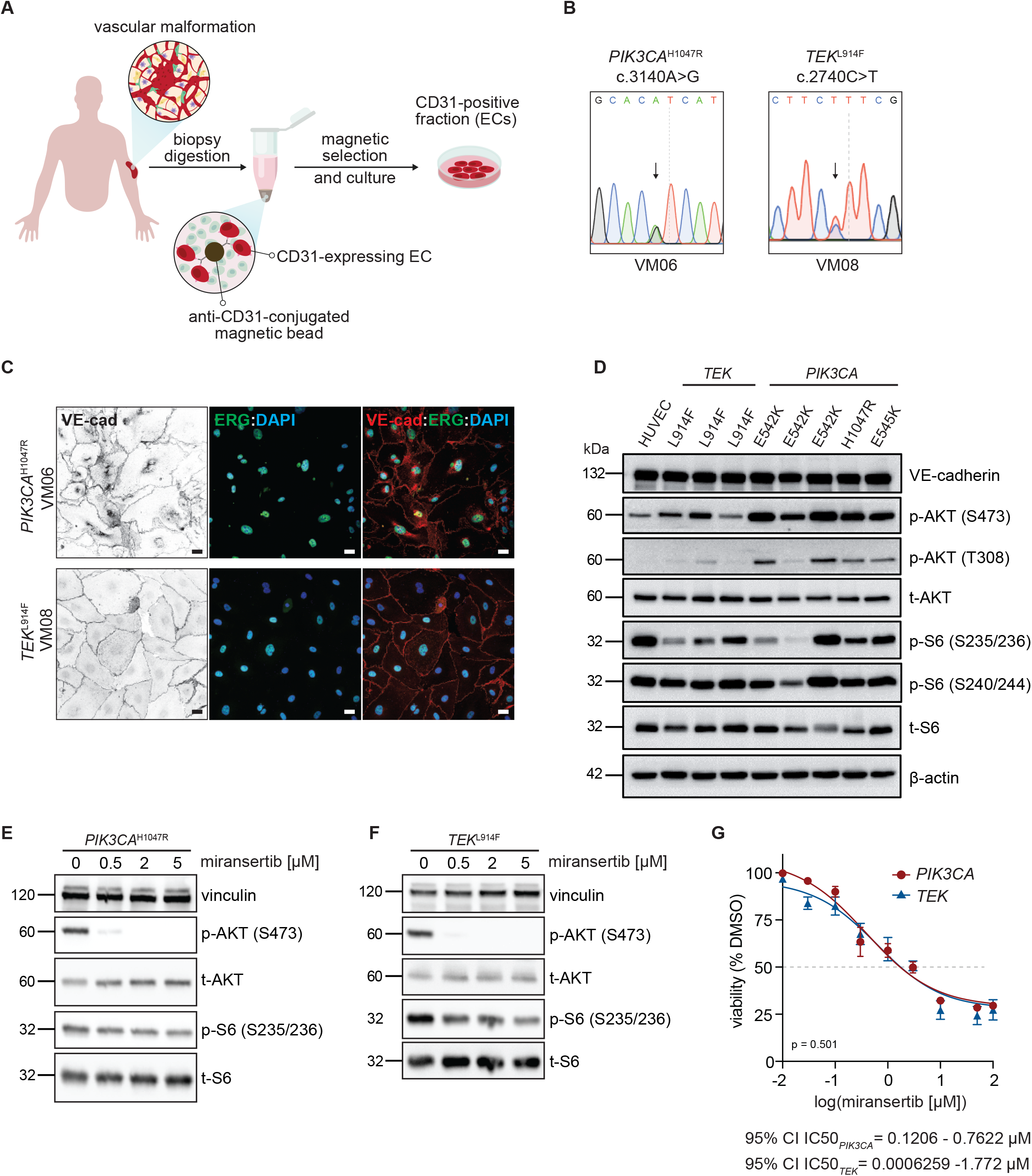
Miransertib impairs cell viability of *PIK3CA*- and *TEK*-mutant patient-derived ECs. (**A**) Illustration showing the strategy of EC isolation from patient-derived biopsies. (**B**) Representative sequencing chromatograms of *PIK3CA* and *TEK* mutant VM-derived ECs. Arrows show the detected point mutations. (**C**) Representative confocal images of *PIK3CA* and *TEK* patient-derived ECs immunostained for VE-cadherin (EC-specific junctional protein) and ERG (EC-specific transcription factor). Cell nuclei were visualised with DAPI. Scale bars: 30 μm. (**D**) Immunoblot showing the activation of PI3K/AKT/mTORC1 pathway (by assessing the levels of p-AKT and p-S6) among different *PIK3CA* and *TEK* patient-derived ECs. Primary HUVEC were used as wild type control. (**E-F**) Immunoblot showing the impact of miransertib at increasing doses on PI3K/AKT/mTORC1 pathway (by assessing p-AKT and p-S6 levels) in (**E**) *PIK3CA* and (**F**) *TEK* mutant patient-derived ECs. (**G**) *PIK3CA* and *TEK* mutant EC viability upon the treatment with miransertib for 72h at different doses assessed by MTS assay. Fitting curves and 95% CI IC50 values for both *PI3KCA* and *TEK* ECs are shown. Statistical analysis was performed by comparison of best-fit values using the extra sum-of-squares F test.

## Discussion

Activation of the PI3K/AKT signalling pathway by genetic mutations in the endothelium is the primary etiological cause of most, if not all, low-flow vascular malformations (Castillo *et al*, 2016a; Castel *et al*, 2016; Limaye *et al*, 2015; Boscolo *et al*, 2015; Luks *et al*, 2015). Despite of this, there is no molecular targeted therapy approved for their clinical management today. Using the postnatal mouse retina as a model of vascular malformations in combination with a tamoxifen-induced strategy which allows to mosaically express the activating H1047R *Pik3ca* mutation at different developmental times, we demostrate that active angiogenesis is required for *Pik3ca* mutants to generate vascular malformations. Here we report on an optimised, robust, and efficient preclinical system displaying traits constituting the main hallmarks of low-flow blood vascular malformations’ pathogenesis: overactivation of PI3K signalling, low-flow vascular compartment specificity, loss of mural cell coverage, and EC hyperproliferation. With this model we show that low dose miransertib effectively target these hallmarks preventing and inducing the regression of already formed vascular malformations driven by *Pik3ca* overactivation.

Low-flow vascular malformations are congenital diseases therefore causative mutations occur during embryonic development (Pang *et al*, 2020). However, the mechanisms of how mutations result in a malformed vascular bed remained elusive. Our results demonstrate that activating *Pik3ca* mutations in ECs are not sufficent for a malformed vascular lesion to appear and that a burst of growth factor signals is essential to generate and expand them. Indeed, we demonstrate that *Pik3ca*-related vascular malformations *in vivo* form via EC hyperproliferation during active angiogenesis. In line with this, cultured *TEK* or *PIK3CA* mutant endothelial cells lead to pathological proliferative response only upon stimulation of growth signals (Cai *et al*, 2019; Le Cras *et al*, 2020). Also, *in vivo*, other types of PI3K-related vascular malformations such as *Pik3ca*-driven lymphatic malformations or hereditary hemorrhagic telangiectasia-like arteriovenous malformations rely on growth factor signals to be induced and expand (Mäkinen *et al*, 2021; Martinez-Corral *et al*, 2020; Garrido-Martin *et al*, 2014). Another important observation of our model is that *Pik3ca*-related blood vascular malformations only occur in veins and in the capillary bed, excluding arteries which remain unaltered. This resembles the human spectrum of vascular malformations where *PIK3CA* mutations have not been reported in arterial malformations. Why arteries are unaffected by *Pik3ca* mutation remains unclear. This could be related to the refractory behaviour that arterial cells exhibit upon reaching their definitive maturation state (Orsenigo *et al*, 2020; Luo *et al*, 2021) and the cell cycle arrested state which fluid shear stress imposes on arterial ECs (Fang *et al*, 2017).

The growth-push concept fits with clinical observations where low-flow vascular malformations appear during embryonic development, expand proportionally with the physiological growth of the patient and are largely quiescent during adulthood, when the production of growth factors is residual (Pang *et al*, 2020). This also explains why low-flow vascular malformations are histopathologically characterized by a low rate of EC proliferation (Castillo *et al*, 2016b). However, despite of being considered as non-proliferative lesions at the time of diagnosis, acute production of extrinsic inputs such as hormonal changes, injury, or wound healing, reactivate malformed vasculature (Pang *et al*, 2020); thereby being a key aspect to mind for preventive approaches. In fact, this may explain why lesions recur upon incomplete surgical removal when wound healing signals are acutely produced. Collectively, our data suggest that blocking EC growth should be considered during potential reactivation scenarios. Furthermore, our study calls for revisiting current classification of low-flow vascular malformation which should be considered proliferative vascular disorders.

Inhibition of the PI3K/AKT signalling pathway has been prioritised in targeted therapy strategies in medical oncology (Vanhaesebroeck *et al*, 2021; Castel *et al*, 2021). However, for vascular malformations there is no molecular therapy approved in the clinic today. This is surprising given that *PIK3CA* oncogenic mutations causing vascular malformations are also present in epithelial cancer (Samuels *et al*, 2004). At least in part, the lack of robust and rapid preclinical models for drug evaluation has hampered the field. In this study, we provide evidence that the mouse retina model offers an opportunity for preclinical drug testing of blood vascular disorders. The key aspects of this model are: (i) it develops in a short time frame (one week); (ii) it allows for a dynamic preclinical testing which covers prevention and curative strategies; (iii) it grants the quantification of disease hallmarks including vascularity, EC proliferation and PI3K signalling, thus providing a robust and non-biased setting to evaluate treatment efficacy; (iv) it recapitulates relevant features of vascular malformations microenvironment including the impact of blood flow, extracellular matrix, surrounding pericytes, and smooth muscle cells, which may influence drug response. Of relevance, this experimental system has been previously used for targeted therapy validation in other vascular disorders (Ola *et al*, 2018, 2016; Alsina-Sanchis *et al*, 2018). Importantly, our model takes into account the mosaic and focal nature of the disease, in which vascular malformations arise in an otherwise normal vasculature network. Thus, preclinical analysis in isolated and focal vascular malformations embedded in a normal vascular plexus allows the study of the impact of drugs in both the lesion and the normal vasculature. This permits the identification of potential knock-on effects of compounds in the systemic vasculature. All in all, we propose mouse retinas as the standard pre-clinical model for low-flow blood vascular malformations.

Blocking AKT signalling has been a promising therapy for cancer; however, rewiring PI3K signalling due to the complex genetic landscape and instability of malignant cells have dampened these expectations (Jansen *et al*, 2016). In contrast, vascular malformations are considered monogenic diseases caused by activating point mutations in an otherwise stable genetic context (Castel *et al*, 2020). These mutations cause an overactivation of the PI3K pathway by constitutive phosphorylation of wild type AKT. Miransertib is a pan-AKT allosteric inibitor and has been shown as a promising AKT inhibitor in preclinical and clinical studies (Lazaro *et al*, 2020) by inhibiting overactive wild-type AKT (Kostaras *et al*, 2020). This therapeutic approach has proven to be efficent in preclinical models of cancer (Yu *et al*, 2017) and in Proteus syndrome, caused by the mosaic expression of an activating *AKT1* mutation (Biesecker *et al*, 2020; Lindhurst *et al*, 2015). Of relevance, miransertib has shown clinical benefit in two children with PROS (Forde *et al*, 2021). Altogether, it has led to the current phase 1/2 clinical trial (NCT03094832) for the assessment of miransertib in Proteus and PROS patients. Based on this, we aimed at assessing the impact of miransentib as a novel treatment for isolated low-flow blood vascular malformations. This seems particulary relevant for those patients in which PI3K inhibitors cause severe side effects due to the pleiotropic functions of PI3K. Here, we provide the first *in vivo* demonstration of miransertib efficacy in PI3K-driven vascular malformations. Our data indicate that, similar to alpelisib, a selective p110α inhibitor, low dose miransertib is sufficent to prevent and induce complete regression of these diseases (Venot *et al*, 2018). We anticipate that these results will have a direct impact on future clinical strategies. Indeed, these patients may need long-term or even chronic treatment; thus an effective low dose may turn specially relevant to avoid undesirable side effects for paediatric age patients.

To complement our *in vivo* preclinical approach, we have set up the isolation and culture of human primary ECs from low-flow vascular lesions. On average, these cultures showed a 50% allelic frequency of *PIK3CA-* or *TEK*-mutant allelels which is suggestive of clonal culture and expansion. Of note, all human cells exhibited overactivation of AKT compared to non-mutant cells, although with allele differences. This unique material turns really valuable to test the impact of targeted therapies. In this regard, we demonstrate that miransertib similarly impairs cell viability of *PIK3CA-* and *TEK*-mutant ECs derived from patients. This is in line with previous studies reporting the effect of miransertib in *PIK3CA*-mutant endothelial cells (Boscolo *et al*, 2019). Our results might indicate that this therapy is also effective for *TEK*-mutant vascular malformations; however, the lack of *TEK*-mutant mouse lines precludes the assessement *in vivo*.

The discovery that most low-flow lesions are caused by the overactivation of PI3K signalling has catalysed the repurpose of PI3K pathway inhibitors for these diseases. In order to promote their direct clinical application, the field urgently needs reliable preclinical models before and while clinical trials are running in humans. By using mouse retinas, we solve this limitation while providing formal proof that low dose miransertib is sufficent to prevent and treat *Pik3ca*-related vascular malformmations. Complementing these data with primary human ECs offers a unique opportunity for personalized medicine.

## Materials and Methods

### Reagents

All chemical reagents were purchased from Sigma-Aldrich, unless stated otherwise. Cell culture media and buffers were purchased from Lonza and Gibco. Primers were obtained from Invitrogen.

### Mice

The *in vivo* experiments were performed in agreement with the guidelines and legislations of the Catalan Ministry of Agriculture, Livestock, Fisheries and Food (Catalonia, Spain), following protocols approved by the local Ethics Committees of IDIBELL-CEEA. Mice were kept in individually ventilated cages under specific pathogen-free conditions. All mice were crossed onto the C57BL/6J genetic background. *Pik3ca*^WT/H1047R^ mice carry a single, germline Cre-inducible point mutation in *Pik3ca* allele (H1047R) (Kinross *et al*, 2012). This mice are crossed onto *Pdgfb*-iCreER mice (Claxton *et al*, 2008) that express an inducible iCreER recombinase from the endogenous *Pdgfb* locus (EC specific). Control mice were CreiER-negative littermates injected with 4-hydroxytamoxifen. iCreER-mediated recombination in *Pik3ca*^WT/H1047R^ mice was induced by intraperitoneal injection of 4-hydroxytamoxifen (doses indicated in the figure legends). ROSA-mTmG double fluorescent reporter mouse (Muzumdar *et al*, 2007) was crossed to *Pdgfb*-iCreER mice. The ROSA-mTmG allele was kept heterozygous. 4-Hydroxytamoxifen was injected intraperitoneally in indicated doses (see the figure legends) and mouse retinas isolated at indicated time points. Cre-mediated recombination was assessed by the expression of membrane-bound GFP.

### Pharmacological in vivo treatment

Miransertib (ARQ 092·2MSA salt) (ArQule, Inc., a wholly-owned subsidiary of Merck & Co., Inc., Kenilworth, NJ, USA) was prepared at a stock concentration of 10 mgA/ml in 20% Captisol (m/v) in 0.02 M citrate/saline buffer. 20% Captisol (m/v) in 0.02 M citrate/saline buffer was used as a vehicle. Mice were injected intraperitoneally with either 75 mg/kg or 35 mg/kg dose of miransertib.

### Mouse retina isolation and immunostaining

Mice were sacrificed by decapitation and eyes were isolated, followed by an hour incubation on ice in 4% PFA in PBS. Isolated retinas were fixed for additional hour, permeabilised overnight at 4°C in permabilisation/blocking buffer (1% BSA, 0.3% Triton X-100 in PBS). Afterwards, the retinas were incubated overnight at 4°C with specific primary antibodies, diluted in permabilisation/blocking buffer (ERG (Abcam, AB92513, diluted 1:400), NG2 (Milipore, AB5320, diluted 1:200), p-S6 235/236 (Cell Signalling Technology, 4857, diluted 1:100). Samples were washed three times in PBS contraining 1% Tween-20 (PBST), following incubation with PBlec buffer (1% Triton X-100, 1 mM CaCl_2_, 1 mM MgCl_2_ and 1 mM MnCl_2_ in PBS, pH 6.8) for 30 minutes at RT. Secondary AlexaFluor-conjugated antibodies, diluted in PBlec, were added to the retinas and incubated for another 2 hours (Invitrogen, A11001, A11011, A11008, A31573). Blood vessels were visualised with AlexaFluor-conjugated Isolectin GS-B4 (Molecular Probes, I21411, I21412). Following three washes with PBST, the tissues were flat-mounted on a microscope slide.

### In vivo proliferation assay

5-ethynyl-2’-deoxyuridine (EdU)-incorporation assay has been performed using a commercially available kit (Invitrogen, C10340). Animals were injected intraperitoneally with 60 µl of EdU (0.5 mg/ml in 50% DMSO and 50% PBS solution) and after 2 hours the animals were sacrificed and retinas isolated. EdU-incorporation was detected with Click-iT EdU Alexa Fluor-647 Imaging Kit, following manufactures instructions. Afterwards, standard protocol for retina immunostaining was applied.

### Confocal imaging and image quantification

Microscopy imaging was done with Leica TCS SP5 confocal microscope. Volocity, Adobe Photoshop 2021 and ImageJ softwares were used for image editing and quantification, respectively. Images were taken from at least 4 retina areas in each genotype. At least three biological replicates per genotype were performed. To quantify the vascular lesion area, an IB4-positive area was manually selected and the percentage of IB4 area per retina area was quantifed. To determine the recombination efficiency of mTmG allele in ECs, the ratio of GFP-positive area to IB4-positive area was calculated and presented as percentage. Retina vascularity was measured using IB4 channel by adjusting the threshold to select the IB4-positive area, followed by quantification of the percentage of IB4-positive area in the total image area (10^4^ µm^2^). EC number was determined manually based on EC-specific nuclei staining (Erg) in 10^4^ µm^2^ image area. Quantification of EC proliferation was done using EdU and Erg co-immunostaining – both EdU- and Erg-positive ECs were quantified in the 10^4^ µm^2^ image area. The coverage of vessels by NG2-positive pericytes was quantified from both NG2 and IB4 channels by adjusting the threshold and selecting the positive NG2 and IB4 areas, respectively. Then the percentage of NG2 to IB4 ratio was calculated. The vascular-specific p-S6 intensity was measured using both p-S6 and IB4 channels. First, a manual threshold was set to obtain the IB4-positive area and define the region of interest (ROI). Then, the integrated density of p-S6 was measured in IB4-positive areas. The background measurements (mean gray values) were taken from areas in close proximity to the vasculature, but negative for IB4. The corrected total fluorescence (CTF) was calculated based on the following equation: CTF = integrated density – (vascular area × mean gray background value).

### Isolation, culture and sequencing of endothelial cells from patient-derived vascular malformations

Patient tissue samples were obtained under therapeutic surgical resection from participants after informed consent with approval of the Committees on Biomedical Investigation at Hospital Sant Joan de Deu and Hospital Santa Creu i Sant Pau (Barcelona, Spain). Collected data were stored in a secure database maintained by Hospital Sant Joan de Deu. Human ECs were isolated from patient-derived biopsies of vascular malformations. Briefly, the biopsy was homogenized with a scalpel blade and digested in dispase II (4 U/ml) and collagenase A (0.9 mg/ml) in Hank’s Balanced Salt Solution for maximum 1.5 hour at 37°C, vortexing the sample every 30 minutes. The digested tissue was disintegrated by pipetting into a single-cell solution, following enzyme inactivation with DMEM supplemented with 10% FBS and 1% penicillin/streptomycin. Cells were resuspended in 100 µl of 0.5% BSA in PBS and incubated with mouse anti-human CD31 (Agilent Dako, M0823, clone JC70A) antibody-coated magnetic beads (ThermoFisher Scientific, 11041) for 1 hour at room temperature. CD31-positive fraction was washed with 0.5% BSA in PBS and sorted with a magnet. Cells were resuspended and cultured in 0.5% gelatin-coated culture well (12-well format) in EGM2 medium (PromoCell, C30140) supplemented with 10% FBS, 1% penicillin/streptomycin (later referred to as EGM2 complete) at 37°Cand 5% CO_2_ until they reach confluency. Cells were subjected for a second selection. Genomic DNA was isolated according to the manufacturer’s protocol (Thermo Fisher Scientific, K182001). The regions of interest in the genomic DNA were amplified by PCR using Platinum™ Taq DNA Polymerase High Fidelity (Thermo Fisher Scientific, 11304011). Exons 10 and 21 of *PIK3CA*, and exon 17 of *TEK* were amplified. PCR products were purified according to the manufacture’s protocol (GE Healthcare, 28-9034-70) followed by Sanger sequencing. Sequences of the primers used: *PIK3CA* exon 10: (forward) 5’-TGGTTCTTTCCTGTCTCTGAAAA-3’ and (reverse) 5’-CCATTTTAGCACTTACCTGTGAC-3’. *PIK3CA* exon 21: (forward) 5’-CATTTGCTCCAAACTGACCA-3’ and (reverse) 5’-TGTGTGGAAGATCCAATCCA-3’. *TEK* exon 17: (forward) 5’-TAGGCAATTTCCACAGCACA-3’ and (reverse) 5’-GGCAAACCAGGCTAAGAGAG-3’. Droplet digital PCR was done on genomic DNA extracted from cell cultures. *PIK3CA* genotyping assays from Bio-Rad were used to specifically detect the *PIK3CA*^*E542K*^ mutation on DNA samples. The Bio-Rad QX200 ddPCR system was used and allelic frequencies were calculated using Quantasoft Analysis Pro (BioRad) software.

### Cell immunofluorescence

Human VM-derived ECs were seeded on gelatin-coated coverslips in a way to reach confluency the next day and incubated overnight at 37°C in 5% CO_2_. Then, cells were washed with warm PBS with Mg^2+^ and Ca^2+^ then fixed with 4% PFA for 15 minutes at room temperature, followed by triple wash with PBS with Mg^2+^ and Ca^2+^. Cells were permeabilised with PBS containing 0.4% Triton X-100 for 5 minutes and blocked with 2% BSA in PBS for 1 hour at RT. The following primary antibodies were used for 1h at RT: VE-cadherin F8 (Santa Cruz, SC-9989, 1:100), ERG (Abcam, ab92513, 1:400). Then, coverslips were washed three times with PBS for 5 minutes, followed by 45 min incubation at RT with appropriate secondary antibody in PBS: goat anti-mouse Alexa Fluor-488 (Invitrogen A11001, diluted 1:300) and goat anti-rabbit Alexa Fluor-568 (Invitrogen A11011, diluted 1:300). Then, coverslips were washed with PBS three times for 5 min, and in the last wash DAPI (Invitrogen, D1306, diluted 1:10 000) was added to visualize cell nuclei. Coverslips were mounted on a microscope slide in a mounting medium (ThermoFisher Scientific, 9990402).

### MTS viability assay and calculation of IC50

MTS assay was used in order to determine cell viability. Briefly, 2·10^3^ cells were seeded on gelatin-coated 96-well plates (5 technical replicates per condition) and incubated overnight at 37°Cin 5% CO_2_ atmosphere. The next day, cells were treated for 3 days with miransertib. MTS assay (Abcam, ab197010) was performed for 2.5 hours and the absorbance was measured at 490 nm. Data (the percentage of a vehicle) were plotted against the logarithm of inhibitor concentration. IC50 and 95% CI values were calculated by non-linear regression (variable slope) using GraphPad Prism software.

### Protein extraction and immunoblotting

Cells were lysed in ice cold lysis buffer (50 mM Tris-HCl pH 7.4, 5 mM EDTA, 150 mM NaCl and 1% Triton X-100) containing protease (Roche, 11836153001) and phosphatase inhibitors (Sigma-Aldrich, 4906837001). Total cell lysates were resolved on 10% polyacrylamide gels, transferred onto nitrocellulose membranes and incubated with appropriate primary and secondary antibodies. The following primary antibodies were used: p-AKT S473 (CST, 4060, diluted 1:1000), p-AKT T308 (CST, 4056S, diluted 1:500), AKT (CST, 9272, diluted 1:2000), p-S6 S235/236 (CST, 4857, diluted 1:1000), p-S6 S240/244 (CST, 2215S, diluted 1:1000), S6 (CST, 2212, diluted 1:1000), VE-cadherin (Santa Cruz Biotechnology, sc-6458, diluted 1:500), vinculin (Abcam, ab49900, diluted 1:10000). The following secondary antibodies from DAKO were used (all diluted 1:5000): swine anti-rabbit (P0399), rabbit anti-goat (P0449), rabbit anti-mouse (P0260), and rabbit anti-sheep (P0163).

### Statistics

Statistical analysis was performed by a nonparametric Mann Whitney’s test using Prism 8 (GraphPad Software Inc.) unless indicated otherwise. All figures are displayed with individual data points that indicate biological replicates and with the standard error of the mean (s.e.m.) as errors bars. At least 3 biological replicates were used. P values considered as statistically significant were as follows: *p < 0.05; **p < 0.01 and ***p < 0.0001.

## Acknowledgements

We thank members of the Endothelial Pathobiology and Microenvironment Group for helpful discussions. This work was partially founded by ArQule, Inc., a wholly-owned subsidiary of Merck & Co., Inc., Kenilworth, NJ, USA. We thank CERCA Program/Generalitat de Catalunya and the Josep Carreras Foundation for institutional support. M.G. laboratory is supported by the research grants SAF2017-89116R-P (FEDER/EU) from MCIU (Spain) co-funded by European Regional Developmental Fund (ERDF), a Way to Build Europe; PTEN RESEARCH Foundation (BRR-17-001); La Caixa Foundation (HR18-00120; also to E.B. and J.M.); by la Asociación Española contra el Cancer (AECC)-Grupos Traslacionales (GCTRA18006CARR); by la Fundación BBVA (Ayuda Fundación BBVA a Equipos de Investigación Científica 2019); World Cancer Research (21-0159). Personal support was from Marie-Curie ITN Actions (P.K. and J.Z.) grant agreement 675392.

S.D.C. is a recipient of a fellowship from the European Union’s Horizon 2020 Research and Innovation Programme under the Marie Sklodowska-Curie grant agreement No 749731. S.D.C. is currently funded by la Caixa Banking Foundation Junior Leader project (LCF/BQ/PR20/11770002). E.B. is funded by the Agencia Estatal de Investigación (Proyectos de investigación en salud PI20/00102). The authors thank the Xarxa de Bancs de Tumors de Catalunya (XBTC; sponsored by Pla Director d’Oncologia de Catalunya). We are grateful to the Band of Parents at Hospital Sant Joan de Déu for supporting the overall research activities of the Developmental Tumor laboratory (Pediatric Cancer Center Barcelona).

## Author contribution

M.G., S.D.C., P.K. and H.S. were the main contributors in the conception, design, acquisition, and interpretation of the data and in writing the article. P.K. H.S, O.V., A.A-U., L.M., J.Z., N.G. O., C.L. and S.D.C. performed experiments and data analysis with input from S.D.C. and M.G.. C.R. and O.M-A. interpreted histopathology. V.C., S.L., E.B and J.M. liaised with human subjects and provided access to human tissue samples and clinical input in the study.

## Conflict of interest

M.G. has a research agreement with ArQule, Inc., a wholly-owned subsidiary of Merck & Co., Inc., Kenilworth, NJ, USA, and Venthera. E.B. is founder and CAB of Venthera; PI and Advisor for Pierre Fabre; PI of the clinical trial NCT04589650 (Novartis).

## The Paper Explained

PROBLEM: Low-flow vascular malformations are congenital diseases caused by a focal overgrowth of vessels. They may cause pain, bleeding, infections, and obstruction of organs. Despite that their genetic causes are known for some years, which lead to the hyperactivation of PI3K signalling, at present there is no molecular targeted therapy approved for these diseases. RESULTS: We have generated a robust and fast preclinical *in vivo* model that allows for testing of targeted drugs. With this model we have demonstrated that PI3K-driven vascular malformations rely on active angiogenesis to occur. Our preclinical studies show that AKT inhibition using low dose miransertib prevents the disease and fully regresses established vascular malformations.

IMPACT: Our new *in vivo* model of PI3K-driven vascular malformations is a reliable and fast preclinical setting to test new or repurposed targeted drugs. Our studies support that *Pik3ca*-mutant endothelial cells cause vascular malformations upon growth stimuli highlighting the importance of preventive therapeutic approaches after invasive treatments. We provide proof of concept for the use of the AKT inhibitor miransertib in PI3K-driven vascular malformations which opens a new window for targeted therapeutic intervention for these diseases. Also, we demonstrate *in vitro* that this targeted therapy is similarly effective in both *PIK3CA* and *TEK*-mutant vascular malformations.

