## Supplemental Figures for "Low dose AKT inhibitor miransertib cures PI3K-related vascular malformations in preclinical models of human disease"

### Figure EV3

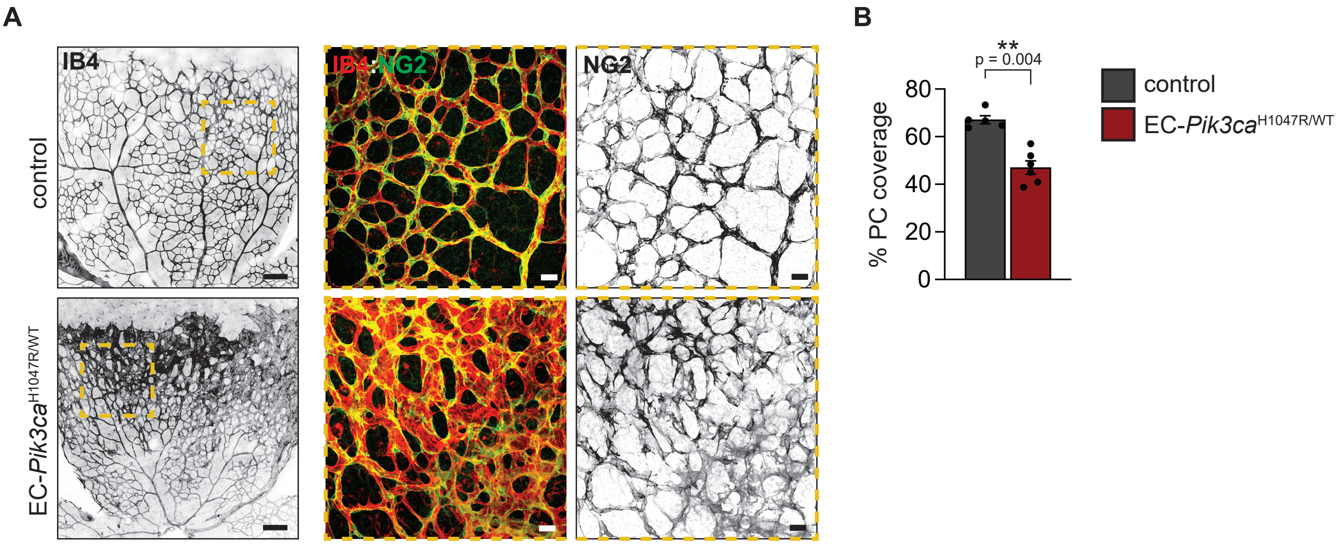

A
IB4
IB4 NG2
NG2
control
EC-Pik3ca H1047R/WT
B
**
p = 0.004
% PC coverage
80
60
40
20
0
control
EC-Pik3ca H1047R/WT

**Figure EV4**

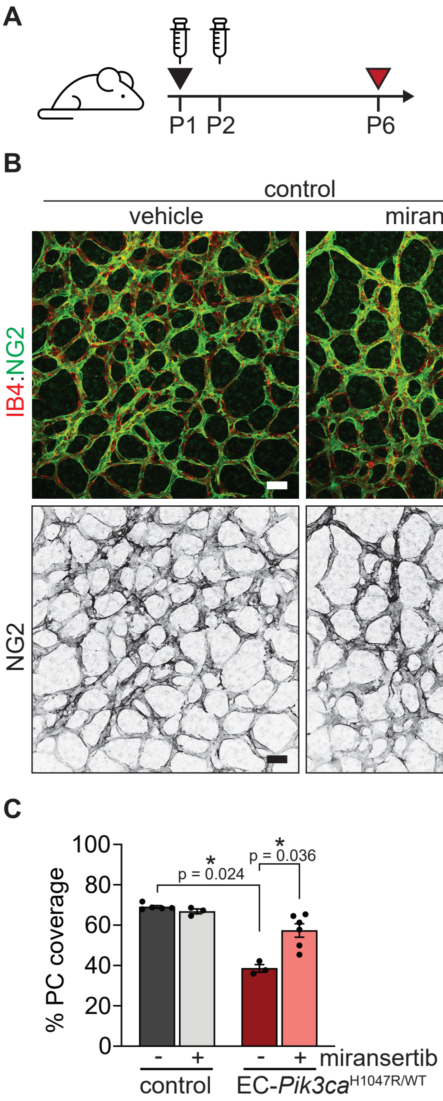
A
0.125 mg/kg 4-OHT
75 mg/kg miransertib/vehicle
retina isolation
P1 P2
P6
B
control
EC-Pik3ca H1047R/WT
vehicle
miransertib
vehicle
miransertib
IB4:NG2
NG2
C
% PC coverage
100
80
60
40
20
0
p = 0.024
p = 0.036
*
*
-
+
-
+
miransertib
control
EC-Pik3ca H1047R/WT

### Figure EV6

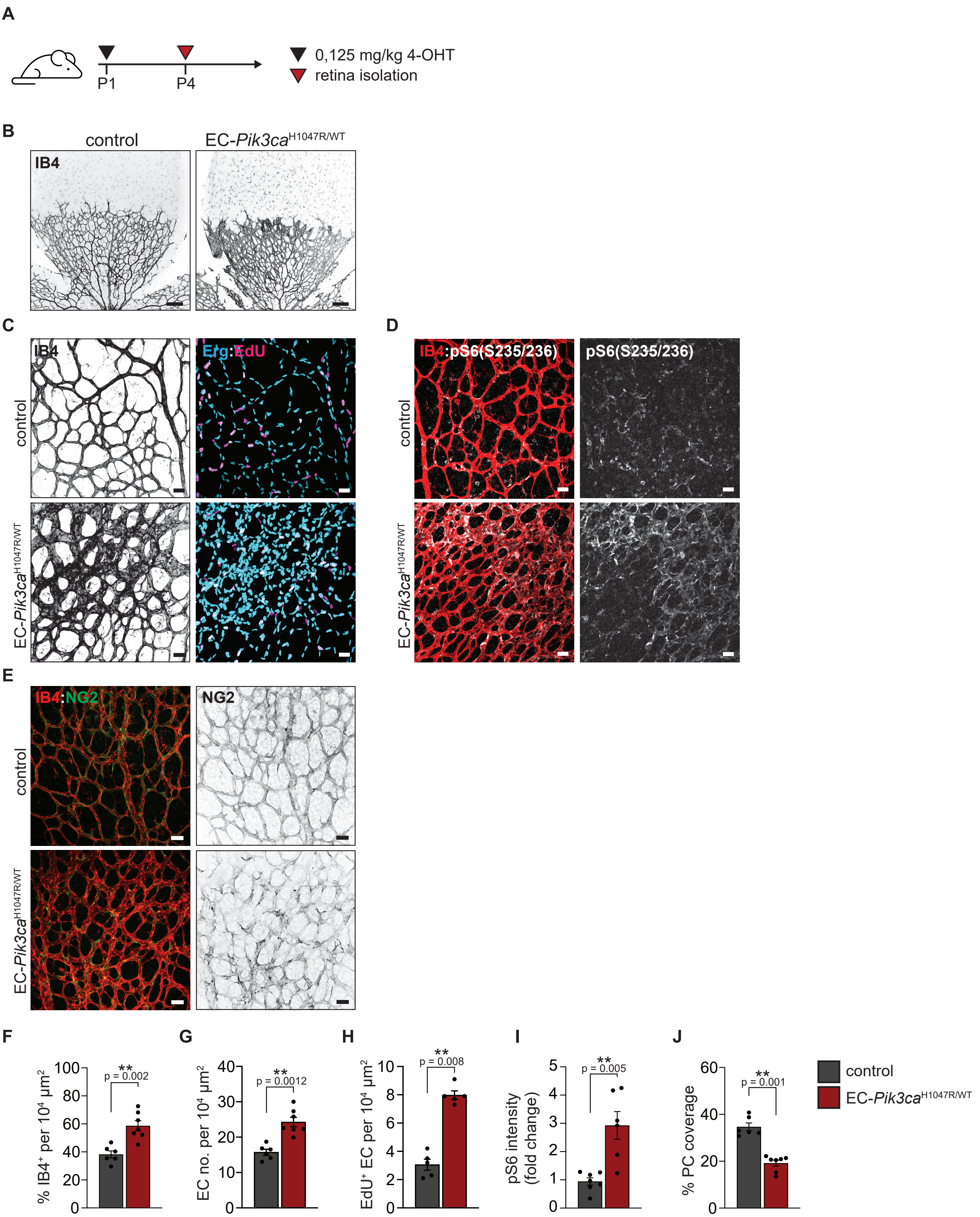

A
0,125 mg/kg 4-OHT
retina isolation
P1
P4
B
control
EC-Pik3caH1047R/WT
IB4
C
IB4
Erg:EdU
control
EC-Pik3caH1047R/WT
D
IB4:pS6(S235/236)
pS6(S235/236)
E
IB4:NG2
NG2
F
% IB4+ per 10^4 µm^2
p = 0.002
G
EC no. per 10^4 µm^2
p = 0.0012
H
EdU+ EC per 10^4 µm^2
p = 0.008
I
pS6 intensity (fold change)
p = 0.005
J
% PC coverage
p = 0.001
control
EC-Pik3caH1047R/WT

**Appendix table S1**

| ID | Genotype | | Sex | Age at surgery | Type | Location |
|---|---|---|---|---|---|---|
| VM01 | *PIK3CA* | E542K | female | 10 | venous malformation | hand |
| VM02 | *PIK3CA* | E542K | female | 56 | venous malformation | hand |
| VM03 | *PIK3CA* | E542K | male | 2 | venous malformation | nd |
| VM04 | *PIK3CA* | E542K | male | 5 | venous - lymphatic malformation | leg |
| VM05 | *PIK3CA* | E545K | male | 15 | venous - lymphatic malformation | nd |
| VM06 | *PIK3CA* | H1047R | female | 62 | venous malformation | finger |
| VM07 | *TEK* | L914F | female | 38 | venous malformation | lip |
| VM08 | *TEK* | L914F | male | 50 | venous malformation | finger |
| VM09 | *TEK* | L914F | female | 10 | venous malformation | lip |
| VM10 | *TEK* | L914F | nd | 1 | venous malformation | face and neck |
| VM11 | *TEK* | L914F | female | 3 | venous malformation | face |

### Appendix figure S1

**A**

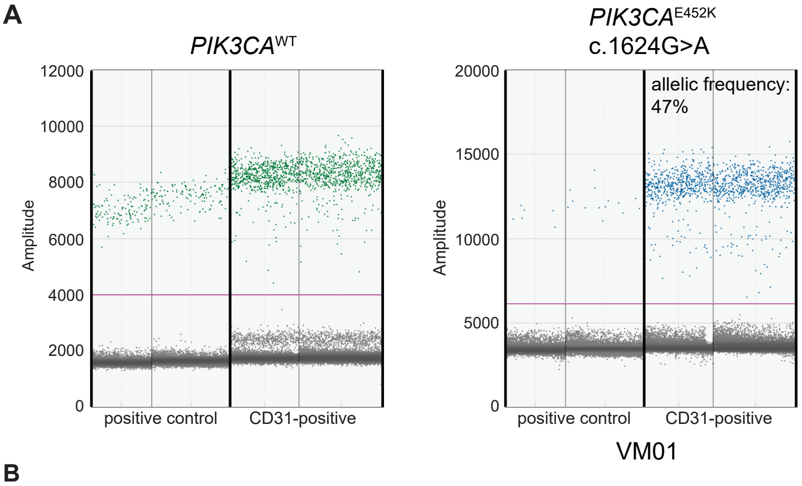


**B**

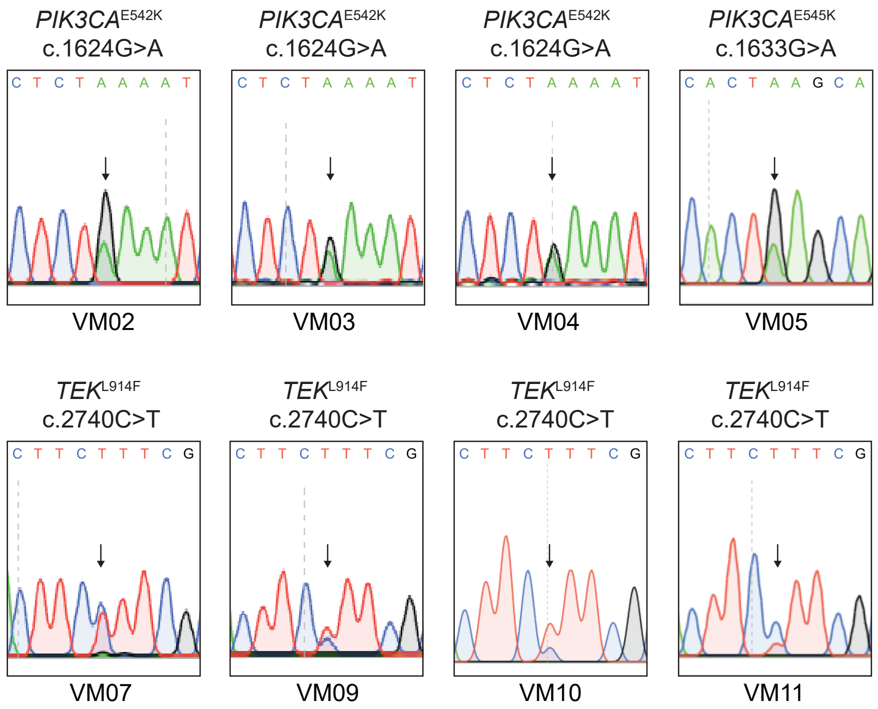


**C**

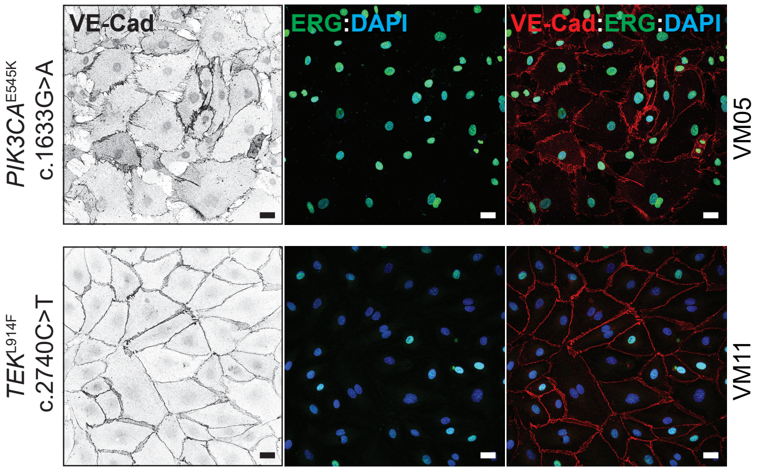

### Expanded View Figure Legends

**Figure EV3. *Pik3ca*-driven vascular malformations exhibit reduced coverage by pericytes.** (**A**) Representative images of control and EC-*Pik3ca*$^{H1047R/WT}$ P6 retinas immunostained for blood vessels (IB4) and pericyte marker (NG2). Dashed areas with high magnifications shown on the right. Scale bars: 150 µm (left panels) and 30 µm (right panels). (**B**) Quantification of vessel coverage by pericytes. Data presented as a percentage of pericyte coverage over EC area (IB4 staining). Error bars are s.e.m. $n \geq 5$ retinas per genotype. Statistical analysis was performed by nonparametric Mann–Whitney test. $^{**}p < 0.01$ was considered statistically significant.

**Figure EV4. Miransertib prevents from loss of pericyte coverage.** (**A**) 4-OHT and miransertib dosing scheme used for a prevention therapeutic experimental setup. (**B**) Representative images of P6 retinas isolated from control and EC-*Pik3ca*$^{H1047R/WT}$ mouse littermates. Blood vessels were stained with IB4 and NG2 immunostaining was used to visualize pericytes. Scale bars: 30 µm. (**C**) Quantification of vessel coverage by pericytes. Data presented as a percentage. Error bars are s.e.m. $n \geq 3$ retinas per genotype. Statistical analysis was performed by nonparametric Mann–Whitney test. $^{*}p < 0.05$ was considered statistically significant.

**Figure EV6. EC-*Pik3ca*$^{H1047R/WT}$ P4 retinas exhibit vascular malformations.** (**A**) Scheme showing 4-OHT treatment regime. (**B**) Representative images of P4 retinas from control and EC-*Pik3ca*$^{H1047R/WT}$ immunostained for blood vessels (IB4). Scale bars: 150 µm. (**C**) Representative high magnification images showing blood vessels (IB4), EC nuclei (Erg) and proliferative cells (EdU). (**D**) Representative high magnification images of retinas immunostained for blood vessels (IB4) and pS6 (S235/236). (**E**) Representative high magnification images showing blood vessels (IB4) and pericytes (NG2). Scale bars (C, D and E) = 30 µm. Quantification of (**F**) retina vascularity, (**G**) EC number, (**H**) EC proliferation by EdU staining, (**I**) pS6 intensity and (**J**) pericyte coverage in control and EC-*Pik3ca*$^{H1047R/WT}$ P4 retinas. Error bars are s.e.m. $n \geq 5$ retinas per genotype. Statistical analysis was performed by nonparametric Mann–Whitney test. $^{**}p < 0.01$ was considered statistically significant.

### Appendix Figure Legends

**Appendix table S1.** Clinical data from patients indicating genotype of vascular malformation, sex of patient, age at surgery, type of vascular malformation and location of vascular malformation. nd, not determined.

**Appendix figure S1. Characterization of primary ECs isolated from patient-derived vascular malformations.** (**A**) Digital Droplet PCR-based detection of the $PIK3CA^{E542K}$ mutation in cultured ECs (CD31-positive) from the vascular malformation of patient VM01 and a positive control. The plots indicate the digital PCR droplets positive with wild type (green), mutant (blue) amplicons. (**B**) Sequencing chromatograms for *PIK3CA* and *TEK* mutant VM-derived ECs. Arrows show the detected point mutations. (**C**) Representative confocal images of *PIK3CA* and *TEK* patient-derived ECs immunostained for VE-cadherin (EC-specific junctional protein) and ERG (EC-specific transcription factor). Cell nuclei were visualised with DAPI. Scale bars: 30 µm.
